# Expression of Lanthipeptides in Human Cells

**DOI:** 10.1101/2023.10.19.563208

**Authors:** Sara M. Eslami, Imran R. Rahman, Wilfred A. van der Donk

## Abstract

Cyclic peptides represent a burgeoning area of interest in therapeutic and biotechnological research. In opposition to their linear counterparts, cyclic peptides, such as certain ribosomally synthesized and post-translationally modified peptides (RiPPs), are more conformationally constrained and less susceptible to proteolytic degradation. The lanthipeptide RiPP cytolysin L forms a covalently enforced helical structure that may be used to disrupt helical interactions at protein-protein interfaces. Herein, an expression system is reported to produce lanthipeptides and structurally diverse cytolysin L derivatives in mammalian cells. Successful targeting of lanthipeptides to the nucleus is demonstrated. In vivo expression and targeting of such peptides in mammalian cells may allow for screening of lanthipeptide inhibitors of native protein-protein interactions.

**Table of contents graphic:** 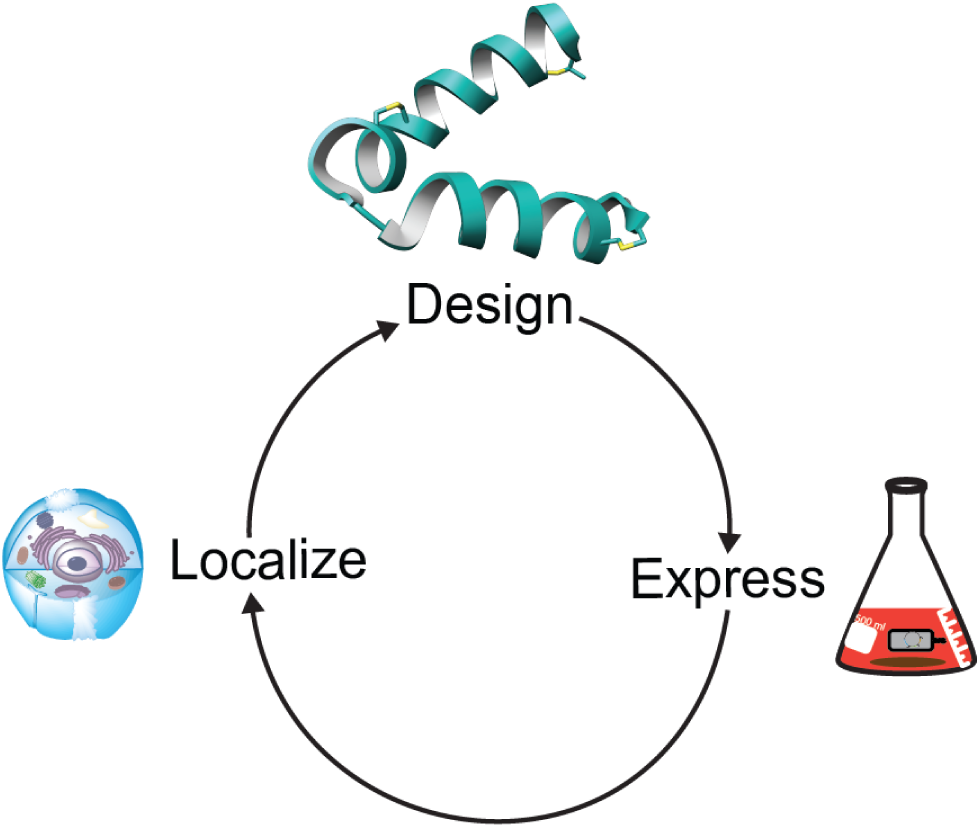

## Introduction

Cyclic peptides represent a burgeoning area of interest in therapeutic and biotechnological research.^1–8^ In opposition to their linear counterparts, cyclic peptides are more conformationally constrained and less susceptible to proteolytic degradation.^9^ Such peptides therefore present unique opportunities in the design of protein-protein interaction (PPI) inhibitors and enhanced or novel bioactivities.^4,6,10–13^ As the catalogue of ribosomally synthesized and post-translationally modified peptides (RiPPs) continues to grow,^14^ an ever-expanding area of research has focused on the biotechnological applications of RiPP structures and pathways.^14,15^

Current methods for the generation of cyclic peptide libraries that can be interrogated for target binding or disrupting PPIs include phage display, bacterial display, yeast display, mRNA display, split-intein circular ligation of peptides and proteins (SICLOPPS), and split-and-pool synthesis.^1,8,10,16–24^ Methods such as mRNA display and split-and-pool synthesis use in vitro peptide screening, making them particularly applicable for extracellular targets. Conversely, in cellulo cyclization using SICLOPPS allows identification of intracellular functional inhibitors of PPIs using monocyclic peptides.^8,25^ Display technologies for enzymatically cyclized, polycyclic peptides have been developed in bacteria,^17^ phage,^18,19^, and yeast^19^ but have also typically been screened against extracellular targets. Finally, while a method for intracellular RiPP production combined with a bacterial reversed two-hybrid screen was successfully developed in *E. coli*,^26^ expression of enzymatically produced, multi-cyclic peptides has not been reported in mammalian cells.^27^ Such technology could be valuable because target proteins would be in their native environment in terms of PPIs and post-translational modifications.^8^

First identified in 1934, the two-component lanthipeptide cytolysin is a toxin produced by *Enterococcus faecalis*, an opportunistic pathogen which, upon infection, may cause endocarditis, urinary tract infections, endophthalmitis, and alcoholic hepatitis.^28–30^ Isolation of the pAD1 plasmid encoding the genes for cytolysin biosynthesis revealed that two peptides, CylL_L_ and CylL_S_ are modified by a single LanM enzyme (CylM), and subsequently cleaved into their mature bioactive forms (CylL_L_’’ and CylL_S_’’) via the consecutive actions of the proteolytic activators CylB and CylA.^31,32^ Structural characterization showed that both peptides are class II lanthipeptides, with CylL_L_’’ containing three (methyl)lanthionines and CylL_S_’’ containing two thioether rings (Figure 1A).^33,34^ With the successful expression of cytolysin in an *E. coli* heterologous host, both CylL_L_’’ and CylL_S_’’ were demonstrated to be amenable to mutagenesis and structure-activity relationship (SAR) studies.^35^ Although cytolysin SAR has been explored, showing that the lanthipeptide synthetase CylM is tolerant towards changes in its substrate sequence, little has been done in pursuit of applying CylL_L_’’ or CylL_S_’’ for biotechnological applications. Given the covalently enforced helical structure of CylL_L_’’ due to three thioether staples as determined by nuclear magnetic resonance (NMR) spectroscopy (Figure 1B),^36^ and the frequent occurrence of helices at the interface in PPIs,^37–40^ CylL_L_’’ may be suitable as a template for disrupting protein-protein interactions.

**Figure 1:**
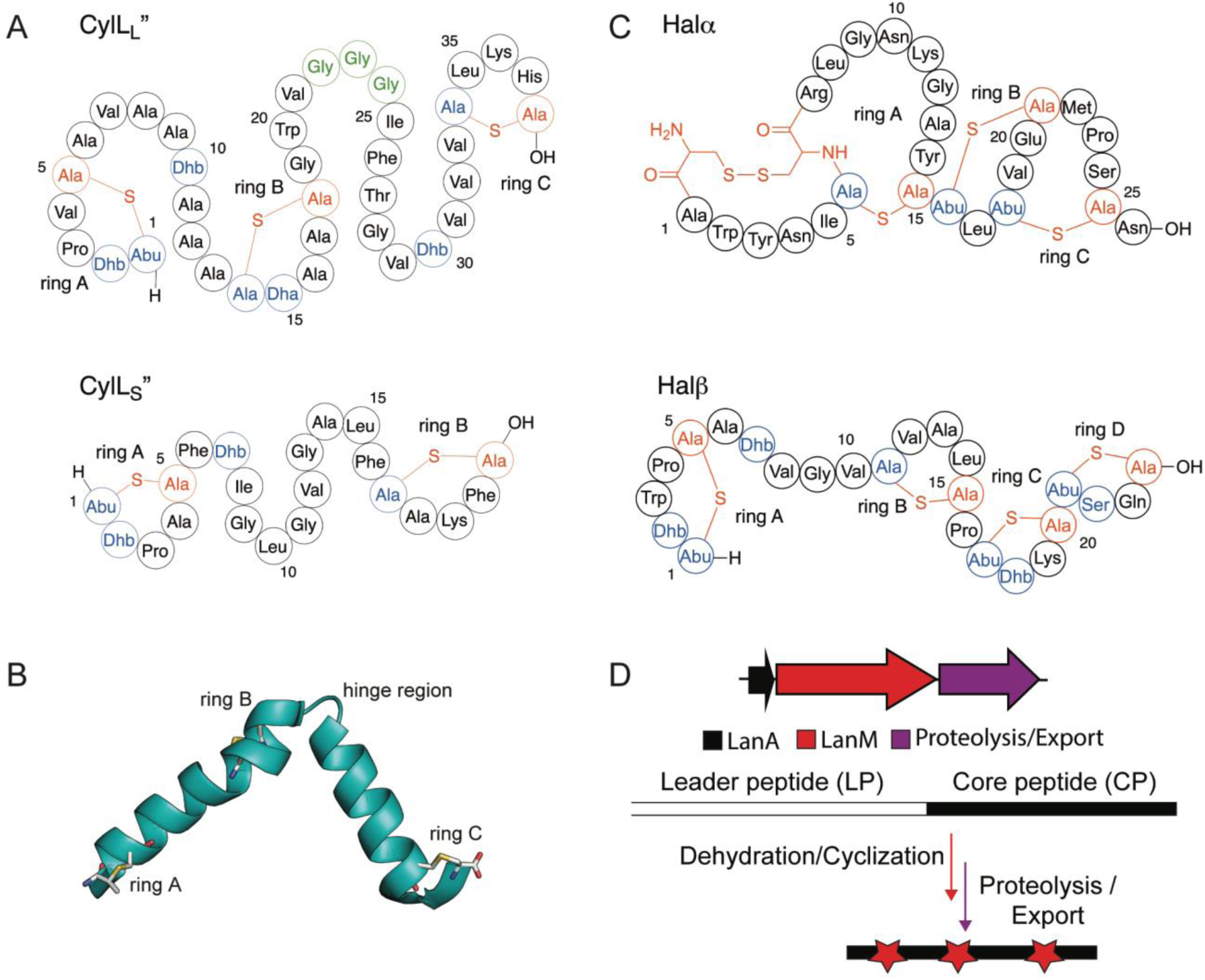
Structure of the two-component cytolysin composed of CylL_L_’’ and CylL_S_’’ (A) and the two-component lanthipeptide haloduracin (B). Dehydrated residues are shown in blue while Cys residues involved in ring formation are shown in red. The hinge region of CylL_L_” is outlined in green. (C) NMR structure of the CylL_L_” subunit and (D) general biosynthetic machinery of class II lanthipeptides.

Class II lanthipeptides are biosynthesized in bacteria as ribosomally synthesized precursor peptides (LanA) with a leader peptide that is important for recognition by the LanM lanthionine synthetases, and a core peptide that is converted to the final mature structure (Figure 1D).^41^ The LanM proteins first dehydrate Ser and Thr residues in the core peptide by phosphorylating the side chain alcohols followed by phosphate elimination to form dehydroalanine (Dha) and dehydrobutyrine (Dhb), respectively.^42^ The same LanM protein then catalyzes Michael-type addition of the thiols of Cys residues to the Dha and Dhb residues.^43^ In the case of cytolysin, the substrate peptides are denoted CylL_L_ and CylL_S_ and the synthetase is CylM.^33^ The leader peptide can be removed with the protease CylA.^44^ In the case of haloduracin β, part of another class II two-component lanthipeptide (Figure 1C), the substrate is HalA2 and the enzyme HalM2.^45^

Herein, we describe the heterologous expression of CylL_L_ and HalA2 with their modification enzymes CylM and HalM2 in mammalian cells leading to the successful installation of multiple macrocyclic thioether linkages. We also show successful localization of the lanthipeptides to the nucleus, demonstrating the potential in nuclear targeted PPI disruption. Our work provides a method for cyclic peptide generation in mammalian cells, illustrating the applicability and utility of this system for future biotechnological applications.

## Results

### CylL_L_’’ adopts a helical secondary structure in a biologically relevant environment

Of the two cytolysin components, the longer CylL_L_’’ forms a helical structure with the first helix comprised of rings A and B connected via a glycine linker (hinge region) to the second helix containing ring C (Figure 1B). The presence of such secondary structures is common within hemolytic peptides that target cell membrane components.^46–53^ Importantly, CylL_L_’’ requires the presence of CylL_S_’’ for cytolytic and antimicrobial activity and requires removal of the leader peptide to be activated. Hence, we did not anticipate production of modified CylL_L_, which still contains the leader peptide, to be toxic to the cell. Previous NMR structural analysis revealed that the two CylL_L_’’ helices can adopt either an extended, linear conformation, or a more compact conformation with the second helix hinged at an angle. The secondary structure of CylL_L_’’ was determined in methanol which may have favored helix formation.^29,36^ To determine if helicity of the peptide is maintained in a physiologically relevant environment, the peptide was dissolved in PBS buffer and analyzed via circular dichroism (CD) spectroscopy. Analysis of the spectrum demonstrated local minima at 208 nm and 222 nm indicative of an alpha helical secondary structure (Figure 2). With this confirmation of helical conformation of the post-translationally modified peptide in buffer, expression of CylL_L_ and CylM was pursued in a HEK293 adherent cell line.

**Figure 2:**
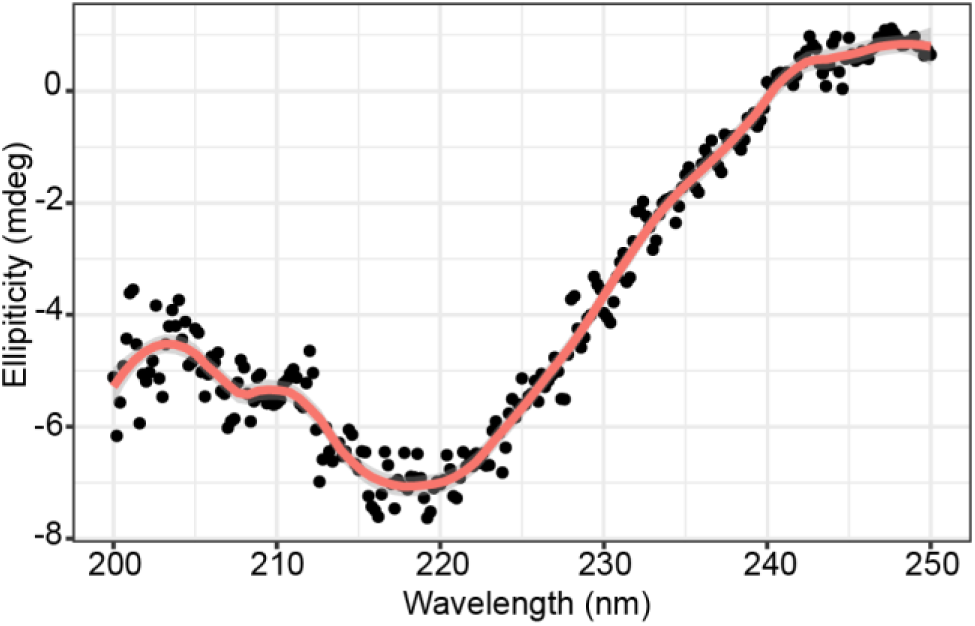
Circular dichroism analysis of CylL_L_” in PBS. Five scans were obtained with PBS as a blank. Wavelengths were monitored from 200 – 250 nm. Data were smoothed with a Savitsky-Golay filter and visualized in R with loess smoothing (span = 0.2).

### Design of a lanthipeptide mammalian expression vector

A multi-cistronic vector under the control of the constitutive CMV promoter was used for expression of CylL_L_ (Figure S1). Due to differential codon usage and biases between bacteria and mammalian cells, the genes encoding the lanthipeptide synthetase CylM and its precursor peptide CylL_L_ were codon optimized for *Homo sapiens* expression and commercially synthesized (see Supporting Information for sequences).^54^ The genes encoding CylM and CylL_L_ were separated by the ribosomal-skipping, self-cleaving P2A peptide; P2A was chosen due to the observed higher expression of the second gene as well as high self-cleavage efficiency of the peptide.^55–57^ To expand the versatility of our construct, we also included a FLAG tag and a Yellow Fluorescence-Activating and absorption-Shifting Tag (Y-FAST) at the N-terminus of the peptide. Y-FAST is a small protein which fluoresces upon binding of its fluorogenic substrate.^58^ While the FLAG-tag enables fixed cell immunofluorescence, the advantage of Y-FAST is the potential for dynamic and reversible live-cell imaging. Finally, an N-terminal hexa-His tag was added to the N-terminus of the peptide to allow for purification via Ni^2+^ affinity chromatography. The construct did not contain a selection marker and hence all data shown are from transient transfection. A second plasmid system was constructed with the same design but containing codon-optimized genes encoding HalA2 and HalM2.

### Expression of lanthipeptides in mammalian cells

After expression, the peptides were purified from the cell lysate by metal affinity chromatography. Initial attempts at production of modified CylL_L_ in HEK293 cells resulted in a fully eight-fold dehydrated peptide with a glutathione adduct (Figure S2). Glutathione (GSH) adducts to dehydroamino acids (Dha/Dhb) have been observed previously during heterologous production of lanthipeptides in *E. coli*.^59–62^ LanC-like proteins (LanCL) have been shown to add GSH to peptides in mammalian cells, and indeed *in vitro* LanCL also adds GSH to CylL_L_.^63^ To determine the location of the GSH adduct produced in HEK293 cells, a chymotrypsin digest was performed followed by matrix assisted laser desorption/ionization-time of flight (MALDI-TOF) mass spectrometry (MS). A resultant 1719 Da fragment was analyzed via LIFT fragmentation and shown to correspond to the non-glutathionylated C-terminal portion of the core peptide (Fig. S2). This observation indicated that the GSH adduct was located in the N-terminal portion of the core peptide. Since GSH adducts are more frequently observed on Dha than on Dhb residues because of the higher reactivity of the former, we hypothesized Dha15 (from dehydration of Ser15) to be glutathionylated. Therefore, Ser15 was mutated to Thr and subsequent expression of CylL_L_-S15T in a suspension cell line derivative of HEK293 (Expi293) demonstrated full modification (eight dehydrations) without GSH adducts (Figure 3; Fig. S3A). Furthermore, assessment of potentially uncyclized Cys residues by reaction with iodoacetamide (IAA)^64^ did not show any adducts indicating full cyclization (Figure S3A).

**Figure 3:**
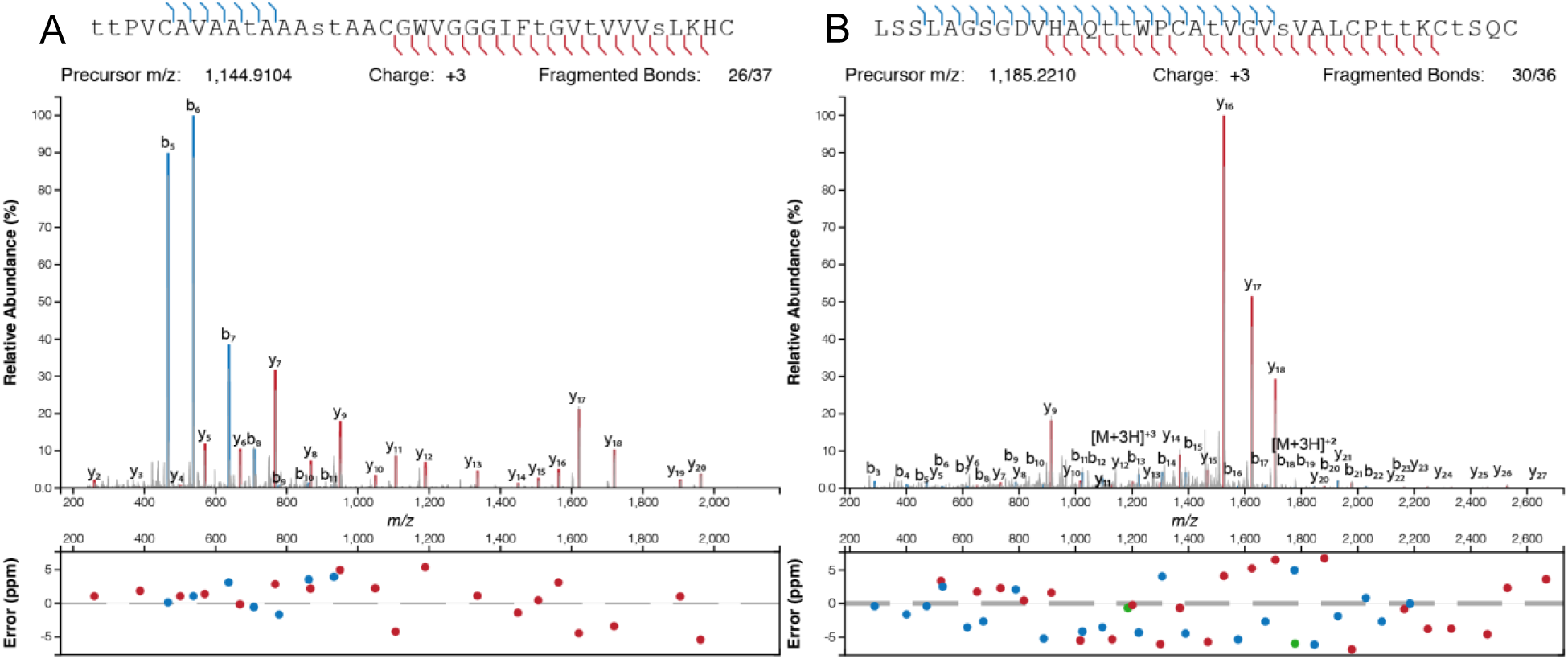
High resolution MS/MS of CylL_L_’’-S15T modified by CylM (A) and HalA2 modified by HalM2 (B) in Expi293 cells. Dehydrated residues are indicated in lower-case. CylL_L_-S15T and HalA2 were digested with CylA and GluC, respectively, prior to HPLC purification and MS analysis.

Following successful production of modified CylL_L_, we applied the method to the production of another class II lanthipeptide, haloduracin β (Halβ). This peptide was chosen as previous research has shown the tolerance of the modifying enzyme HalM2.^19,65^ HalA2 expressed together with the synthetase HalM2 in Expi293 cells was fully dehydrated (Figure 3). Incubation of HalM2-modified HalA2 with N-ethylmaleimide (NEM)^66^ revealed no NEM adducts indicating complete cyclization (Supp. Figure 3B).

### Bioactivity of lanthipeptides produced in mammalian cells

To confirm successful formation of CylL_L_”-S15T and Halβ with correct ring patterns, we tested the peptides produced in Expi293 for bioactivity. CylL_L_” in concert with CylL_S_” (obtained from bacterial expression)^33^ demonstrated robust antimicrobial activity towards Gram-positive bacteria. However, we did not observe growth inhibition when testing the CylL_L_”-S15T mutant against the indicator strain *Lactococcus lactis sp. cremoris*. This observation may be explained by a previous report that mutation of Ser15 severely attenuates anti-microbial activity.^35^ We also assayed Halβ produced in Expi293 for inhibitory activity against the indicator strain *L. lactis* CNRZ 117 when combined with Halα (Figure 1C) produced in *E. coli*.^67^ The α and β subunits alone did not inhibit *L. lactis* but when combined, we observed a zone of inhibition indicating correct structural formation of the β subunit in Expi293 cells (Figure 4).

**Figure 4:**
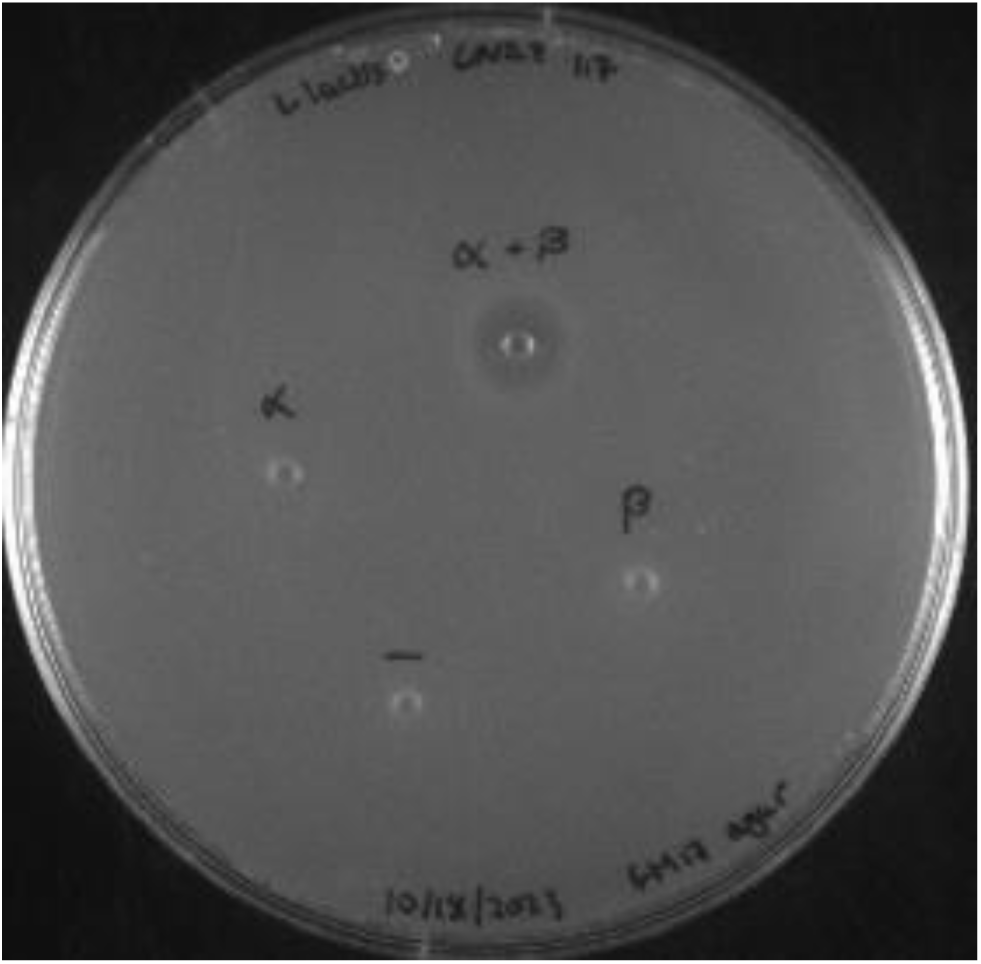
Bioactivity of Halβ produced in mammalian cells. The Halβ core peptide contained two amino acids from the leader peptide (Ala-Gln), but previous studies have demonstrated that even six residues from the leader peptide did not abolish synergistic antimicrobial activity of Halβ when combined with Halα.^68^ Each peptide was tested at 100 pmol individually or combined (α + β) against the indicator strain *L. lactis* CNRZ 117. A deionized H_2_O negative control was included.

### CylL_L_ mutant peptides are expressed and modified in mammalian cells

CylL_L_-S15T was used as a template to generate an NDT (D = A/G/T) mutant library. The five mutated positions were chosen along the same helical face spanning the A and B rings of CylL_L_-S15T (Figure 5). Previous lanthipeptide libraries have employed NNK, NWY, and NDT degenerate codons.^19,26,65^ For this study, NDT codons were chosen for the following reasons. Firstly, it leads to a more diverse array of amino acids including both positively and negatively charged residues (Gly, Val, Leu, Ile, Cys, Ser, Arg, His, Asp, Asn, Phe, Tyr). Secondly, no stop codons are introduced, and thirdly each codon encodes one amino acid thereby preventing overrepresentation of a single amino acid.

**Figure 5:**
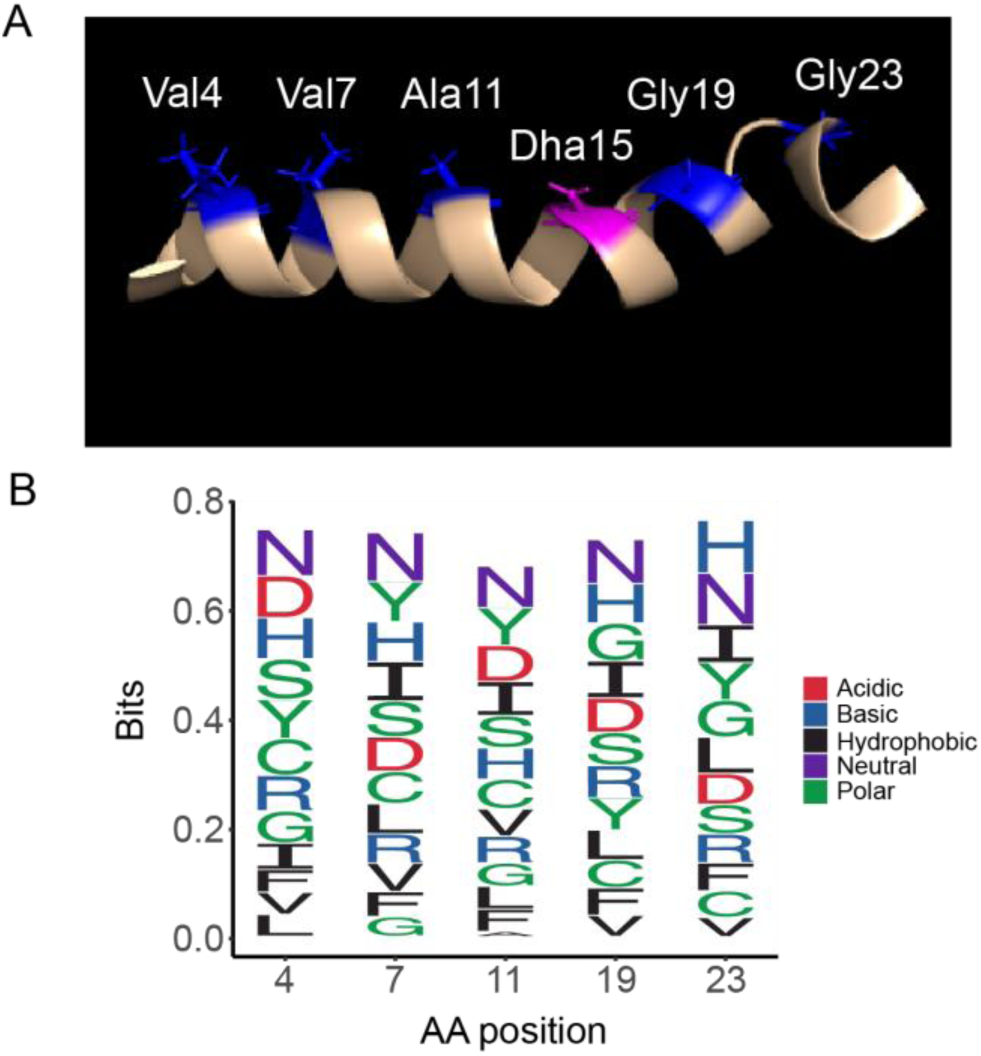
Design of the NDT mutant library of CylL _L_” (A) NMR structure with mutated residues shown in blue and the position of mutated S15T residue in pink. B) Frequency of NDT encoded amino acids at each mutation site in the sequenced library.

Deep sequencing revealed a library size of 1.74 x 10^5^ unique sequences, representing ∼ 70% coverage of the theoretical library size (2.5 x 10^5^). The decreased coverage may be explained by reduced transformation efficiency as well as the observation of a slight overrepresentation of the template sequence during library generation. Generally, however, the distribution of amino acids at the variable positions was near the statistical prediction (Fig. 5B). To determine if CylM could successfully modify CylL_L_-S15T derivatives in mammalian cells, five sequences from the NDT library were chosen randomly for expression in Expi293 cells. Of the five sequences, four were fully modified after co-expression with CylM (Table 1). Dehydrations were confirmed via high resolution tandem mass spectrometry (HR-MSMS) and cyclization was confirmed via iodoacetamide (IAA) assessment of the presence of free thiols (Figure 6). Tandem MSMS data for the four peptides supported similar ring patterns to that of CylL’’ (Supp. Figures 4-7). Mutation of a hinge region residue (Gly23) to Ser in variant 3 did not affect the ring pattern (Supp. Fig. 5).

**Figure 6.**
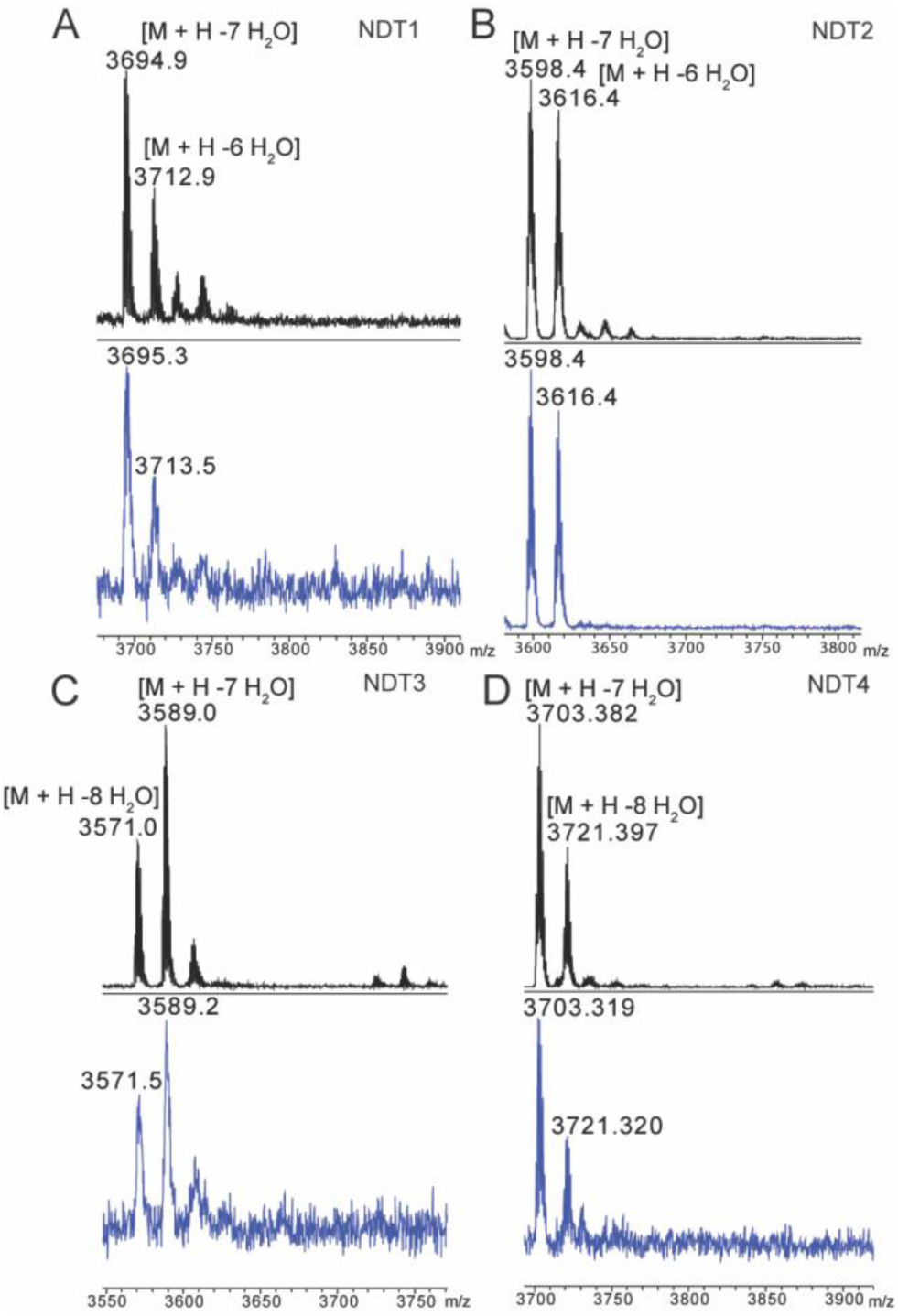
MALDI-TOF mass spectra of CylL_L_-S15T NDT variants modified in Expi293 cells by CylM before (black) and after (blue) iodoacetamide (IAA) reaction. Peptides were expressed in Expi293 cells and purified via Ni-NTA chromatography and analytical HPLC prior to IAA reaction. Reactions were desalted via C4 ziptip.

**Table 1:**
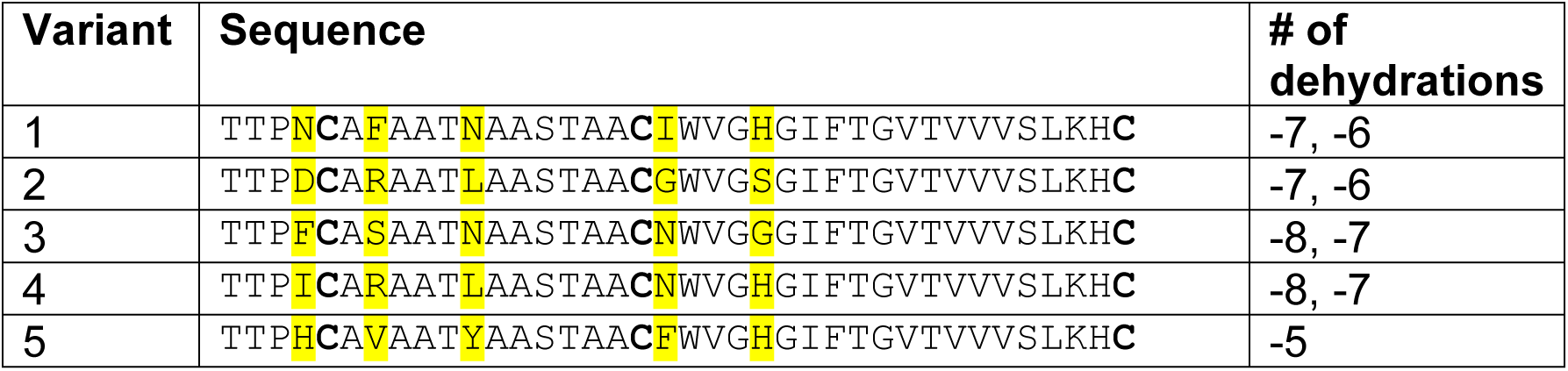
CylL_L_”-S15T NDT variants.

### Lanthipeptides can be targeted to the nucleus in mammalian cells

Given the successful modification ofHalA2 and CylL_L_-S15T and its variants, we next focused on targeted localization of modified CylL_L_-S15T and HalA2 within mammalian cells. Nuclear targeting was chosen because transcription factor interactions are located within the nucleus that may be targets of PPI inhibition. A nuclear localization signal (KKKRKV) was appended to the N-terminus of the FLAG-tagged precursor peptides. Immunostaining of transfected HEK293 cells with an anti-FLAG antibody demonstrated colocalization of CylL_L_-S15T and HalA2 with the nuclear stain DAPI compared to the untargeted controls (Figure 7). To confirm nuclear-targeted lanthipeptides were still fully modified, nuclear localized CylL_L_ and CylM were expressed in Expi293 cells and subsequent purification confirmed full dehydration of the peptide (Supp. Figure 7).

**Figure 7:**
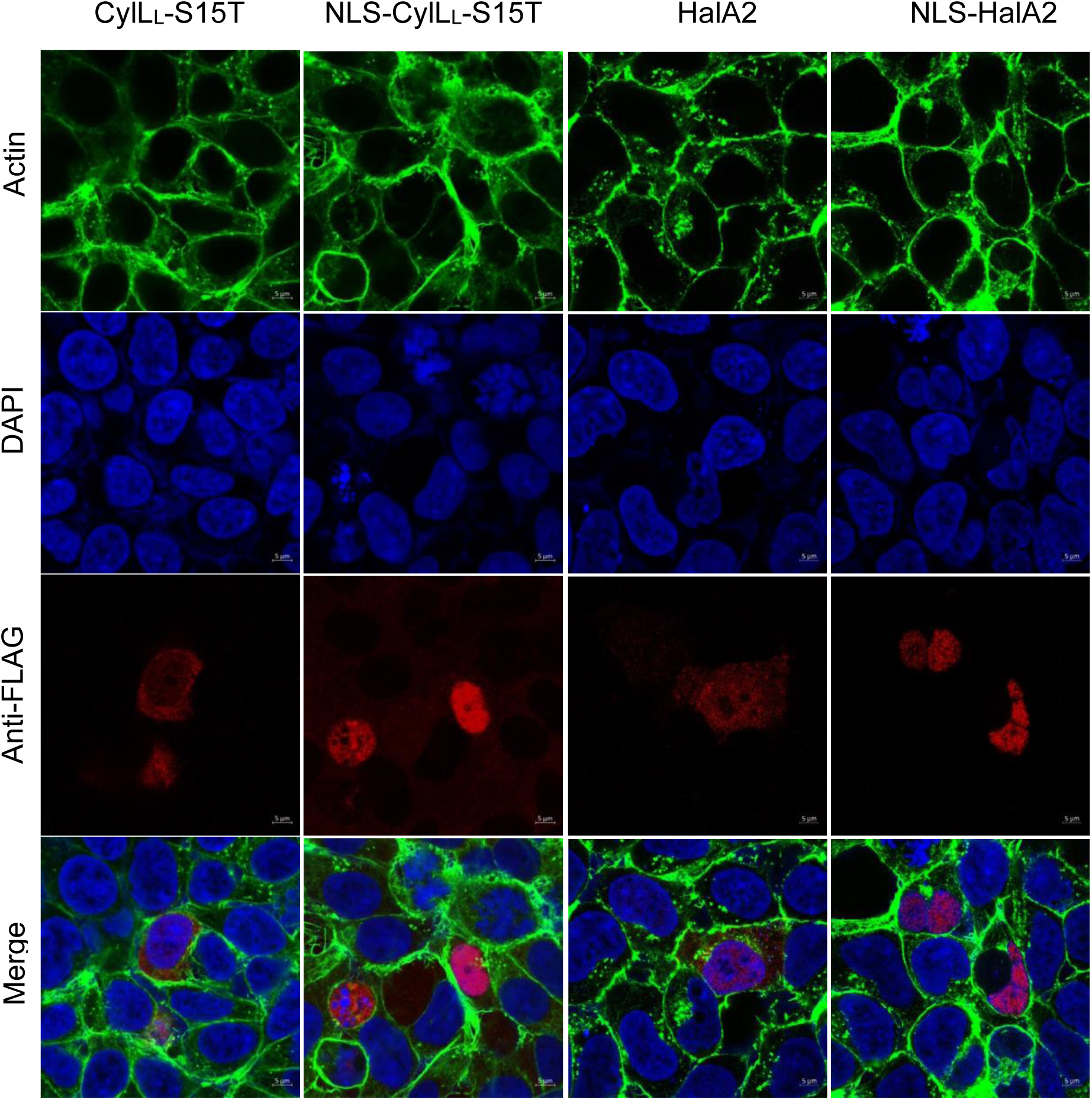
Confocal microscopy of untargeted and nuclear-targeted CylL_L_-S15T and HalA2. HEK293 cells were fixed and permeabilized two days post-transfection. The following stains were used. DAPI (blue, nucleus), phalloidin-488 (green, actin), and mouse Anti-FLAG primary antibody with goat anti-mouse Fluor647 secondary antibody (red, CylL_L_). Immunofluorescence was visualized at 63X magnification using an LSM880 confocal microscope.

## Discussion

Herein we have described a method for the successful production of polycyclic lanthipeptides in mammalian cells as well as the successful nuclear localization of these peptides. Owing to their stability and structural diversity, cyclic peptides have garnered much interest for their therapeutic potential.^4,69^ Previous research has employed display and screening technologies to identify enzymatically generated polycyclic PPI inhibitors in bacteria and yeast.^27^ In vivo expression of such peptides in mammalian cells may allow for functional screening against native protein-protein interactions.

In 2013, the Arora group identified 7,308 helices present at protein interaction interfaces; further analysis revealed over 1,000 helical dimers involved in PPIs of enzymes, transcription factors, and other proteins of interest.^70,71^ Many of these interactions, such as that between MYC-MAX transcription factors, are critical in promoting cell proliferation in tumors; dysregulation of MYC-MAX activity has been shown to increase cancer progression with up to 70% of cancers bearing this hallmark.^71,72^ Disrupting such secondary structures can be accomplished through small molecules, antibodies, or peptides. However, both small molecules and antibodies have limitations in efficacy. Small molecule compounds cannot easily mimic the extended, globular 3-dimensional interfaces and also face high rates of clearance; antibodies, due to their large size, are mainly employed in disrupting extracellular PPIs.^73^ Helical and/or cyclic peptides may fill the gap because of their secondary structures, comparatively small sizes, and resistance to proteolytic cleavage.

Our results demonstrate the utility of lanthipeptide biosynthetic enzymes in the production of a diverse set of polycyclic structures within mammalian cells. Consequently, this method may expand the scope of therapeutic targets accessible to enzymatically generated polycyclic peptides.

## ASSOCIATED CONTENT

Supporting Information. Supporting figures and tables. This material is available free of charge via the Inter-net at http://pubs.acs.org.

## Notes

The authors declare no competing financial interest.

## Acknowledgments

The authors acknowledge the Roy J. Carver Biotechnology Center (CBC) and proteomics core at the University of Illinois at Urbana-Champaign for high resolution mass spectrometry services. We also thank the Core Facilities at the Carl R. Woese Institute for Genomic Biology for the use of a confocal microscope and helpful advice.

## Materials and Methods

### Buffers and media

The pH of all buffers was adjusted using 1 M or 5 M NaOH and HCl and passed through a 0.2 μm filtration device. LanA resuspension buffer (B1) contained 6.0 M guanidine hydrochloride, 0.5 mM imidazole, 500 mM NaCl, 20 mM NaH_2_PO_4_ in DI H_2_O and the pH was adjusted to pH 7.5. LanA wash buffer (B2) contained 20 mM NaH_2_PO_4_, 30 mM imidazole, 300 mM NaCl in DI H_2_O and the pH was adjusted to pH 7.5. LanA elution buffer (EB) contained 20 mM Tris HCl, 1.0 M imidazole, and 100 mM NaCl in DI H_2_O and the pH was adjusted to pH 7.5. LanM start buffer contained 20 mM Tris HCl, 1 M NaCl, pH 7.6 and LanM final buffer was made up of 20 mM HEPES, 300 mM KCl, pH 7.5. A 1X solution of Dulbecco’s phosphate buffered saline (PBS) was used.

### Expression and purification of protease CylA

A culture of 10 mL LB with 10 μL of kanamycin solution (50 mg/mL; Kan50) was inoculated with *E. coli* BL21 containing pRSF_His-CylA^44^ from a frozen stock and incubated overnight at 37 °C, with shaking. A 1 L culture of Terrific Broth (TB) with 1 mL of Kan50 was inoculated with 10 mL of the overnight culture and incubated at 37 °C, with shaking at 220 rpm. The culture was induced with 500 μM IPTG (1 M stock) at mid-log phase and allowed to incubate overnight at 18 °C, with shaking at 200 rpm. After incubation, the culture was pelleted at 4500 *g* for 15 min and the resulting pellet was resuspended in 40 mL of LanM start buffer with lysozyme and allowed to incubate for 20 min on ice. Lysis was performed via sonication (39% amplitude, 4.4 s on, 9.9 s off) for at least 5 min. The lysate was pelleted at 13,000 *g* for 30 min and filtered through a 0.45 μm filter. Purification was performed using immobilized metal affinity chromatography (IMAC) with a 5 mL HisTrap column and an AKTA FPLC. The following conditions were used: the column was equilibrated with 2 – 3 column volumes (CV) of LanM start buffer followed by loading of the lysate; Buffer A = LanA B2, Buffer B = Elution buffer. The column was washed with 10% B for 2 CV, 15% B for 4 CV, and 50% B for 4 CV. Fractions containing CylA were determined via SDS-PAGE with a 4%-20% Tris gel at 200 V and collected, concentrated, and the buffer exchanged into LanM final buffer using a 30 kDa MWCO amicon filter. Aliquots were stored in LanM final buffer with 10% glycerol at -80 °C.

### Expression and purification of CylL_L_ in HEK293 cells

The plasmid construct encoding CylM and CylL_L_ under a constitutive promoter was used to transfect HEK293 cells using Turbofect transfection reagent. HEK293 cells were grown in a 100 mm culture dish and were 70% – 90% confluent at the time of transfection. Cell media was changed 3 – 4 h post-transfection followed by an overnight incubation at 37 °C, 5% CO_2_. After incubation, media was aspirated and the cells were washed with cold 1X PBS. After PBS removal, 5 mL of Pierce IP lysis buffer and ½ of a Pierce protease inhibitor tablet was added to each 100 mm dish and allowed to incubate for 5 – 10 min at 4 °C. The cell lysate was pelleted at 1000 *g* for 5 min and the supernatant was purified via Ni resin affinity chromatography. The supernatant was washed with 3 CV of LanM start buffer with 25 mM imidazole and the peptide eluted in 1 CV of LanM start buffer with 0.5 M imidazole. The elution was treated with 5 – 10 μL of 1.6 μg/μL CylA and allowed to incubate overnight at room temperature. After acidification with 50% TFA/H_2_O, the elution was filtered through a 0.45 μm filter and purified on an Agilent 1260 Infinity analytical HPLC (aHPLC) with a Luna C8 (100 A) analytical column. The following conditions were used with a flow rate of 0.80 mL/min. with solvent A containing H_2_O + 0.1% TFA and solvent B containing ACN + 0.1% TFA: 2% B for 5 min., 2% B – 100% B over 20 min., 100% B for 5 min. UV absorbance was monitored at 220 nm and 254 nm. Fractions were collected manually and analyzed via matrix assisted laser desorption/ionization (MALDI) time-of-flight (TOF) mass spectrometry and LIFT fragmentation using a Bruker UltrafleXtreme.

### Generation of CylL_L_-S15T

Site-directed mutagenesis was performed using overlapping primers. The following polymerase chain reaction (PCR) conditions were used with Phusion polymerase in a 50 μL reaction volume. Each reaction contained 1X Phusion HF buffer, 1 M betaine, 0.2 mM dNTPs, 0.5 μM of each primer, 0.5 μL polymerase, at least 10 ng of DNA template, and DI H_2_O added up to 50 μL. Initial denaturation was performed for 2 min at 95 °C, followed by 26 cycles of denaturation (20 s, 95 °C), annealing (30 s, 69.8 °C), and elongation (7 min., 72 °C), and a final elongation step (72 °C) for 10 min. Successful amplification was confirmed via gel electrophoresis. After amplification, 1 μL of Dpn1 enzyme was added to each 50 μL reaction and incubated at 37 °C for 1 h, followed by a Qiagen PCR cleanup per the manufacturer’s instructions. Subsequently, 5 μL of amplified product was added to 50 μL of chemically competent NEBTurbo cells. Cells were kept on ice for 30 min., followed by heat shock at 42 °C for 30 s and recovery with 1 mL LB for 1 h. After recovery, cells were pelleted and ∼ 700 μL of supernatant was removed; the pellet was resuspended in the remaining liquid, of which 50 μL was plated onto LB/Amp100. Following overnight incubation at 37 °C, single colonies were used to inoculate 10 mL LB/Amp100 and grown overnight at 37 °C. Plasmid extraction (Qiagen miniprep) was performed on the overnight cultures and plasmids were sequenced to determine successful mutagenesis.

### Expression and purification of lanthipeptides in Expi293 cells

Expi293 cells were transfected with expression plasmids per the manufacturer’s instructions. Expression enhancers 1 and 2 were added, per manufacturer’s instructions, 19 – 21 h post-transfection followed by 4 days of incubation at 37 °C, 8% CO_2_. Cells were pelleted at 1000 *g* for 5 min and the resulting supernatant was removed. To each cell pellet, 3 – 5 mL of Pierce IP lysis buffer was added and allowed to incubate with gentle agitation at room temperature for 5 min. The lysate was pelleted at 1000 *g* for 5 min and the supernatant was purified via Ni-NTA affinity chromatography and analytical HPLC as described above.

### Generation of an CylL_L_-S15T NDT mutant library

A primer containing degenerate NDT codons at five amino acid positions along CylL_L_-S15T was used to create a mutant library. The following polymerase chain reaction (PCR) conditions were used with Phusion polymerase in a 50 μL reaction volume aliquoted into 9 μL. Each reaction contained 1X Phusion HF buffer, 1 M betaine, 0.2 mM dNTPs, 0.125 μM of each primer, 0.5 μL polymerase, at least 10 ng of DNA template, and DI H_2_O added up to 50 μL. Initial denaturation was performed for 2 min at 95 °C, followed by 26 cycles of denaturation (20 s, 95 °C), annealing (30 s, 58 °C), and elongation (30 s, 72 °C), and a final elongation step (72 °C) for 5 min. Successful amplification was confirmed via gel electrophoresis. After amplification, 1 μL of Dpn1 enzyme was added to each 50 μL reaction and incubated at 37 °C for 1 h, followed by a Qiagen PCR cleanup per the manufacturer’s instructions. PCR linearized vector and NDT library inserts were assembled via Gibson Assembly (GA) per the manufacturer’s instructions and subsequently dialyzed against DI H_2_O using 0.02 μm membrane. ElectroMAX™ DH10B cells were transformed with the dialyzed GA reactions (2 reactions, 20 μL each) and recovered with 880 μL of Super Optimal broth with Catabolite repression (SOC) media for 1.5 – 2 h. Post-recovery, 100 μL of transformants were plated on Amp100 at 10^-3^ and 10^-4^ dilutions and incubated overnight at 37 °C. The remaining cells were used to inoculate 9 mL of LB (100 μg/mL Amp), incubated overnight at 37 °C, and the library DNA isolated using a Qiagen miniprep kit. NextGen sequencing was performed via SeqCenter and the reads analyzed using a custom Python script.

### Circular dichroism (CD) spectroscopy

CylL_L_’’ was resuspended to 14.8 μM in 1X PBS. Samples were loaded into a 2 mm, Quartz spectrophotometer cell (Starna Cells, cat. no. 1-Q-2) and measured using an Olis Cary-16 circular dichroism spectrometer. Wavelengths in the range of 200 nm – 250 nm were monitored. Five scans were obtained and averaged for each sample. The data were smoothed with a Savitsky-Golay correction in RStudio.

### High-resolution tandem mass spectrometry

Dried peptide samples were suspended in 0.1% formic acid/5% acetonitrile and injected into an UltiMate 3000 rsnLC coupled to a Q Exactive HF-X mass spectrometer (Thermo). Peptides were separated using a 25 cm Acclaim PepMap 100 C18 column (2 µm particle size, 75 µm ID) over the course of 45 min using mobile phases of 0.1% formic acid (A) and 0.1% formic acid in 80% acetonitrile (B); the gradient ran from 5% B to 60% B at 300 nL/min, followed by column washing and re-equilibration. Throughout the course of the run, the column was maintained at 50 °C. The mass spectrometer was operated in the positive mode in a data-dependent manner, where full MS1 scans from 350 to 2000 *m/z* were acquired at 120k resolution (3e6 AGC, 60 ms IT). MS2 scans of the top 15 most abundant ions were acquired at 45k resolution after HCD fragmentation (30 NCE; 5e4 AGC, 35 ms IT). The isolation width was 1.0 *m/z*, and the dynamic exclusion window was 15 s.

AGC: automatic gain control

IT: max injection time

NCE: normalized collision energy

### Bioactivity assay of haloduracin

Modified core peptides were resuspended in sterile, DI H_2_O. The concentrations were then estimated using Pierce Quantitative Colorimetric Peptide Assay according to the manufacturer’s protocol. Agar well diffusion assays were used to evaluate the antimicrobial activity of the modified peptide cores against *L. lactis* CNRZ 117. A starter culture of indicator strain was grown static in M17 medium supplemented with 0.5% glucose under aerobic conditions at 30 °C overnight. Agar plates were prepared by combining molten M17 agar (cooled to 42 °C) supplemented with 0.5% glucose and 300 μL of overnight bacterial culture to yield a final volume of 40 mL. The seeded agar was poured into a sterile Petri dish and allowed to solidify at room temperature after which 100 pmol of modified peptides were spotted onto the agar plates. Plates were incubated at 30 °C overnight, and antimicrobial activity was qualitatively determined by the presence or absence of a zone of growth inhibition.

### Iodoacetamide (IAA) reaction conditions

Purified CylM-modified CylL_L_-S15T mutants were resuspended in 100 μL of DI H_2_O and sonicated in a water bath for at least 5 min. To a 50 μL reaction, 5 μL of 100 mM TCEP, 5 μL of 200 mM KH_2_PO_4_ buffer pH 7.5, and 30 μL of peptide were added. The reaction was incubated for 30 min at 37 °C to allow for reduction of disulfide bonds, prior to addition of 5 μL of 100 mM IAA (in KH_2_PO_4_). Following IAA addition, the reaction was incubated for 30 min at 37 °C. Reactions were purified with C4 ziptip and analyzed via MALDI-TOF MS.

### N-ethylmaleimide (NEM) assay

To a 50 μL reaction, 25 μL of DI H_2_O, 5 μL of 100 mM TCEP, 5 μL of 1 M citrate buffer (100 mM EDTA, pH 6), and 10 μL of peptide were added; it was assumed 100 μg of peptide was present when resuspended in DI H_2_O to a concentration of 100 μM. The reaction was incubated for 10 - 30 min at 37 °C to allow for reduction of disulfide bonds, prior to addition of 5 μL of 100 mM NEM (dissolved in ethanol). Following NEM addition, the reaction was incubated for 30 min at 37 °C. Reactions were purified with C4 ziptip and analyzed via MALDI-TOF MS.

### Confocal microscopy

All steps were performed at room temperature. Cell media was removed and the cells were washed twice with 1 mL of 1X PBS. Cells were fixed in 4% paraformaldehyde for 10 min and washed three times with 1 mL 1X PBS each time. Cells were then permeabilized with 0.5% Triton X-100 (in 1X PBS) for 10 min and washed three times with 1 mL 1X PBS each time. Cells were blocked with 2% bovine serum albumin (BSA) for 1 h and washed two-three times with 1X PBS. Primary anti-FLAG antibody (8146T, Cell Signaling Technologies) diluted 1:1000 in 1X PBS-T and incubated for 3 h. After primary antibody removal and washing (3x with 1 mL 1X PBS), goat anti-mouse-Fluor647 secondary antibody (A-21235, ThermoFisher) was diluted 1:1000 in 1X PBS-T and incubated for 1 h. Following another a wash step (3x with 500 μL 1X PBS), the coverslips were placed onto a slide containing ∼ 7 μL of mounting media containing DAPI. The coverslip was allowed to cure for 30 min and the edges were sealed onto the slide with clear nail polish. Slides were stored at – 20 °C until visualization. Immunofluorescence was visualized using a Zeiss LSM880 microscope with 63X magnification and immersion oil (Immersol^TM^ 518F). Images were analyzed using Zen software (2.3 SP1).

