## Supporting Figures for "Expression of Lanthipeptides in Human Cells"

### Codon optimized primers and genes used in this study

| | |
|---|---|
| SME_CylLL-S15T.F1 | CTGCGTCAACGGCTGCATGCGGCTGGGTGGGCGGCG |
| SME_CylLL-S15T.R1 | GCATGCAGCCGTTGACGCAGCAGCAGTAGCAGCCACG |
| SME_pCMV_NLS.R2 | cactttgcgtttcttcttTGGGCCCGGATTTTCTTCG |
| SME_pCMV_NLS.F2 | aagaagaaacgcaaagtgGACTACAAGGACGACGACGAC |
| SME_NLS_pCMV-FLAG | CGAAGAAAATCCGGGCCCAaagaagaaacgcaaagtgGACTACAAG |
| SME_CylLL_NDT.R3 | ccgtgactccagtgaagattccAHNgcccacccaAHNgcatgcagccgttgacgcagcAHNagtagcagcAHNggcgcaAHNt<br>ggagtcgtc |
| *H. sapiens* codon-optimized CylM | tcagaggacaacctcatcaacgttctttcaatcaacgagagatgtttccttttgaagcaatctgggaaggagaagtacgacataaagaacttg<br>caagcttggaaggagagaaagagtgtgcttaagcaagacgacttggactaccttatcaagtacaagtacgagtctttggacaacttcggact<br>tgggatcacacccatcgagaacttccctgacaaggaagtcgcaatccaatacatcaaggaccaatcctggtacatattcttcgagtcaatact<br>cgactcatacaacgactctgaggagaagctcctcgaggtggacgcttcttaccctttccgctacttcttgcaatacgctcgattgttcttgctcga<br>cctcaactccgagctcaacatctgtactaaagagttcatcataaacctcttggagactctcactcaagagttgatacaccttacatccaagact<br>cttgtccttgacttgcacacgttcaagaagaacgagccactcaaggggaacgacagttccaagagattcatctactacctcaagaagcgatt<br>caactccaagaaggacatcatcgctttctacacgtgttaccccgagctcatgcggatcacagtggtgcggatgagatactttcttgacaatac<br>gaagcagatgctcatccgggtcactgaggacttgccttccatccaaaactgtttcaacatccaatccagcgagctcaacagtatcagtgagtc<br>tcaaggggactcacactccaggggaaagacagtgtccacgctcactttctcagacggcaagaagatcgtctacaagcctaagataaactc<br>cgagaacaagctcagagacttcttcgagttcttgaacaaggagctcgaggcagacatctacatcgttaagaaggttacacgtaacacttactt<br>ctacgaggagtacatagacaatatcgagattaacaatatagaagaggtcaagaagtactatgagcggtacgggaagttgatcggtatcgc<br>gttcctcttcaacgtcactgatctccactacgagaatattatcgcgcacggggagtaccctgtcatcatcgacaacgagactttcttccaacaa<br>aacatacccatcgagttcggcaactctgccactgtcgacgctaagtataagtacttggacagtatcatggtaactgggctcgtgccctaccttg<br>ccatgaaggacaagtccgacagtaaggacgagggtgtgaacctctctgctcttaacttcaaggagcaaagtgtccccttcaagatattgaag<br>atcaagaacactttcactgacgagatgaggttcgagtaccaaacacacatcatggacacagctaagaacacacctataatgaacaacga<br>gaagatttcattcatttcatacgagaagtacatcgttacaggtatgaagtccatcctcatgaaggctaaggactctaaaaaaaagatcctcgc<br>atacatcaacaacaatctccaaaacttgatcgtaagaaacgtcatccgccccacacaacggtacgcggacatgctcgagttcagttaccac<br>ccaaactgtttctctaacgcaatcgagcgggagaaggtccttcacaacatgtgggcttacccttacaagaacaagaaggttgtccactacga<br>gttctctgaccttatagacggagacatccccatcttctacaacaacatctcaaagacatccctcatcgcttccgacggatgcttggtcgaggac<br>ttctaccaagagagcgcattgaaccgatgtttgaacaagataaacgacctttgcgacgaggacatcagtatccaaacagtttggctcgagat<br>agctctcaacatctacaatccatataagtacataaacgaccttaagaatcaaaactccaataagtacatctacactgggctcgagctcaacg<br>gaaagatcatacaagcttgtcaaaagatcgagaagaagattttcaagcgggcaattttcaataagaagacgaacactgtaaactggatcg<br>acataaagcttgaccaagactggaacgttggaatcctcaacaacaacatgtacgacggtctcccaggcatattcatcttctacgtcgcactca<br>agtacatcacaaagaatcacaagtacgactacgtcatagagtgtatcaagaactcaatatacactatcccttccgaggacatcctctccgcct<br>tcttcgggaaggggtccttgatatacccattgttggtggactaccgactcaacaacgacataaacagtctcaacgttgcagtcgagatcgctg<br>acatgctcatagagaagaagcctataaacaacggggagcttaagaacgactggatccacggacacaactccatcatcaaggtcctgcttc<br>ttttgtccgagatcactgaggacgagaagtacaggaagttctcactcgagatcttcgagaagctctccgaggagccctacttcaacttcaggg |

| | |
|---|---|
| | gcttcggacacgggatctactcatacgtccacctcttgtccaagttcaacaggatcgacaaggctaactccttgcttcacaagatcaaggagt<br>catacttcgaagaggagccaaagaacaactcttggtgcaagggaactgtgggcgagctcttggctacgatcgagttgtacgacgacaaca<br>tatccaatattgacatcaataagactatagcgtacaagaacaaggactgtctttgtcacggaaacgctggaacactcgagggcttgatacaa<br>ctcgctaagaaggaccctgagacttaccaatacaagaagaacaagctcatatcatacatgctcaacgacttcgagaagaacaacacact<br>caaggtagcggggagtgagtacttggagtcacttggattcttcgtgggaatctccggagtcggatacgagttgctcagaaaccttgactccga<br>gatccctaacgccttgcttttcgagctc |
| *H. sapiens* codon-optimized CylLL-S15T | tcacaagaccctaactctgagaacctcagtgtagtgccctctttcgaggagctttctgttgaggagatggaagccattcaaggaagtggggat<br>gtgcaagctgagacgactccagtgtgcgccgtggctgctactgctgctgcgtcaacggctgcatgcggctgggtgggcggcggaatcttcac<br>tggagtcacggtagtggtgtccctcaagcactgttga |
| *H. sapiens* codon-optimized HalM | aaaaccccactgacgtcagagcatccttcagtccctacgactctgcctcacactaatgacaccgattggctcgaacaacttcatgacatactc<br>tccatccccgtgactgaggaaatccagaaatattttcacgcagagaacgatctcttctcattcttctatacccctttcctgcaattcacatatcaga<br>gtatgagcgactactttatgacctttaaaaccgatatggccctgattgaacgacaatctcttctccaatctactttaactgctgtacatcatcggct<br>gttccaccttacacatcgcacgctgatctcagagatgcacattgataaacttacagtgggtcttaacggtagcacccctcacgaaaggtatat<br>ggattttaatcacaaattcaacaaaacatccaaatctaagaatctcttcaacatatacccgatactgggcaagctggtagtcaacgagactct<br>ccgaacgataaattttgtaaagaaaattatacaacactatatgaaggactatctcttgctcagtgacttcttcaaggagaaagacctgagactg<br>accaaccttcaactgggagtcggcgacacccatgttaatggccaatgcgtgacaattctgacatttgcgagcggacaaaaggtagtgtataa<br>gccgcgaagtctgtctatcgataagcaatttggcgaatttatagagtgggtaaattcaaagggctttcaaccttctcttcgcattccaatagccatt<br>gacaggcaaacctacggatggtacgagtttataccccatcaagaagctacaagcgaggacgaaattgagcgctattactcccgcataggt<br>ggatacctcgcaattgcttatctcttcggggctacggatctgcacctggataacctcatcgcgtgtggagaacaccccatgctgattgatctcga<br>aaccctcttcactaatgacctggactgctatgattccgcctttccatttccagccttggctcgcgagctcacgcaatcagtattcggcaccctcat<br>gttacccatcactatcgcatctggtaagttattggacattgacctttcagccgtcggcggcggaaaaggcgtccaaagcgagaagattaaaa<br>cttgggtcatcgttaatcaaaagacggatgagatgaaattggttgaacaaccctatgtgactgagagcagccagaataaacccacggtcaa<br>cggcaaggaagcaaatataggcaattacattccacatgtgaccgacgggtttcgcaaaatgtaccgattatttcttaatgaaatcgatgaact<br>gatggatcataatgggcctattttcgcattcgagagctgtcaaatccggcacgtgtttagggctactcatgtatatgccaaatttctggaagcttct<br>actcatcctgattacttacaagaaccaactaggcgcaataaactctttgaaagcttctggaatataacatccctcatggcgccttttaagaaaat<br>tgttccacacgagatcgcagaactcgagaatcatgatattccatatttcgttttgacatgtggcggcaccatcgttaaggatggatacggccga<br>gacatagcagatctgttccaatctagttgtatagaaagggtgactcatagacttcagcaactgggatccgaagacgaggcccgacaaatca<br>gatacattaagtccagtctggcaaccctgaccaacggggactggaccccatcacacgaaaagaccccgatgagtccagctagcgcgga<br>ccgcgaagacgggtattttctccgcgaagcccaagcgattggagatgacattcttgctcaactcatttgggaagatgacaggcacgctgcat<br>acctgatcggtgtctcagtcggcatgaatgaggcggtgaccgtatcccctctgactccaggaatctatgatggtacactggggatcgtgctctt<br>ctttgatcaactcgcgcaacaaacgggtgagacgcattatagacatgctgcagacgcattattggaaggcatgttcaaacaattgaagcca<br>gagttgatgccctcaagcgcatacttcggacttgggagtcttttctacggtcttatggtgctcggtctgcaacgatccgacagccacattataca<br>gaaggcctatgaatatcttaagcacttggaagaatgtgtgcagcatgaagagacgcccgatttcgtgtcaggccttagtggcgtgctctatatg<br>cttaccaaaatatatcagttgactaatgagccacgcgtatttgaggtagccaaaaccacagcttcccgacttagtgtcttattagatagcaaac<br>aacccgacaccgtcttgacaggcttgagtcacggcgccgccggttttgccctcgcgctcctcacctatggtactgccgctaacgacgaacaa<br>ctcctcaaacaagggcactcctatctcgtatatgagcgaaatagatttaataagcaagagaataattgggttgaccttcggaaaggaaatgc<br>ctatcaaacattctggtgtcatggtgcccctggtattggaattagcaggctgttactggctcaattttatgatgacgagttgttgcacgaggaactt<br>aacgccgctctgaataagactataagcgatggttttgggcacaaccattctctgtgccacggagatttcggaaacctcgatctgctgttgctcta<br>cgcccaatatacaaataatccagaaccaaaagaactggccaggaagctggccatatcctctatagatcaggctcatacctatggctggaa<br>actcggcctgaatcacagtgaccaattacaaggaatgatgctcggcgtgacaggaataggataccagttacttagacatataaaccctacg<br>gttccctctatattggccctggaactccccagtagtacactgaccgaaaaggaactcagaatccacgaccgggca |
| *H. sapiens* codon-optimized FactorXa-HalA2 | gttaattctaaagatctccgaaatcctgaatttcggaaagctcagggcctgcaattcgtcgacgaggttaacgagaaagaattgtcaagcctg<br>gccgggtctggagatatagaaggacgaacgacatggccctgtgcaacggtcggagtcagcgtcgcgttatgcccaaccactaagtgcac<br>ctctcagtgctag |

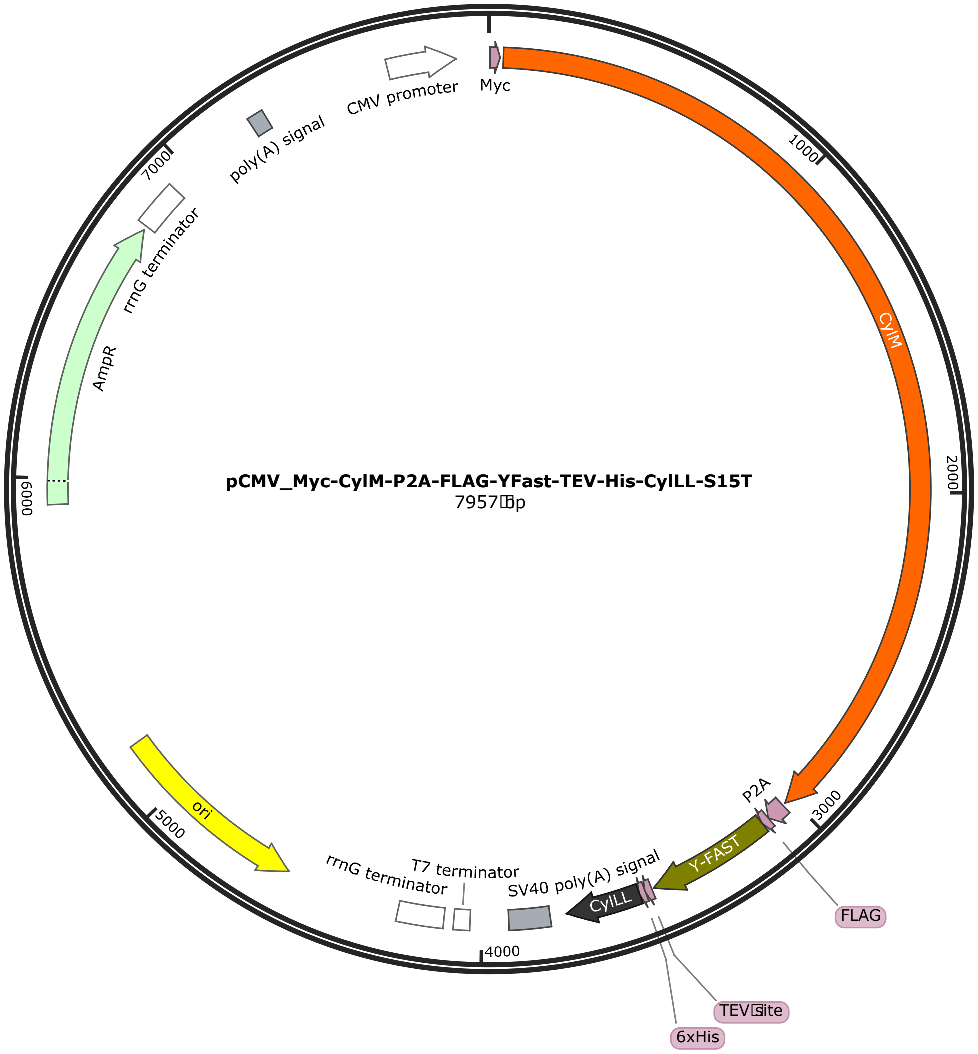


Figure S1: Mammalian expression vector for lanthipeptide production.

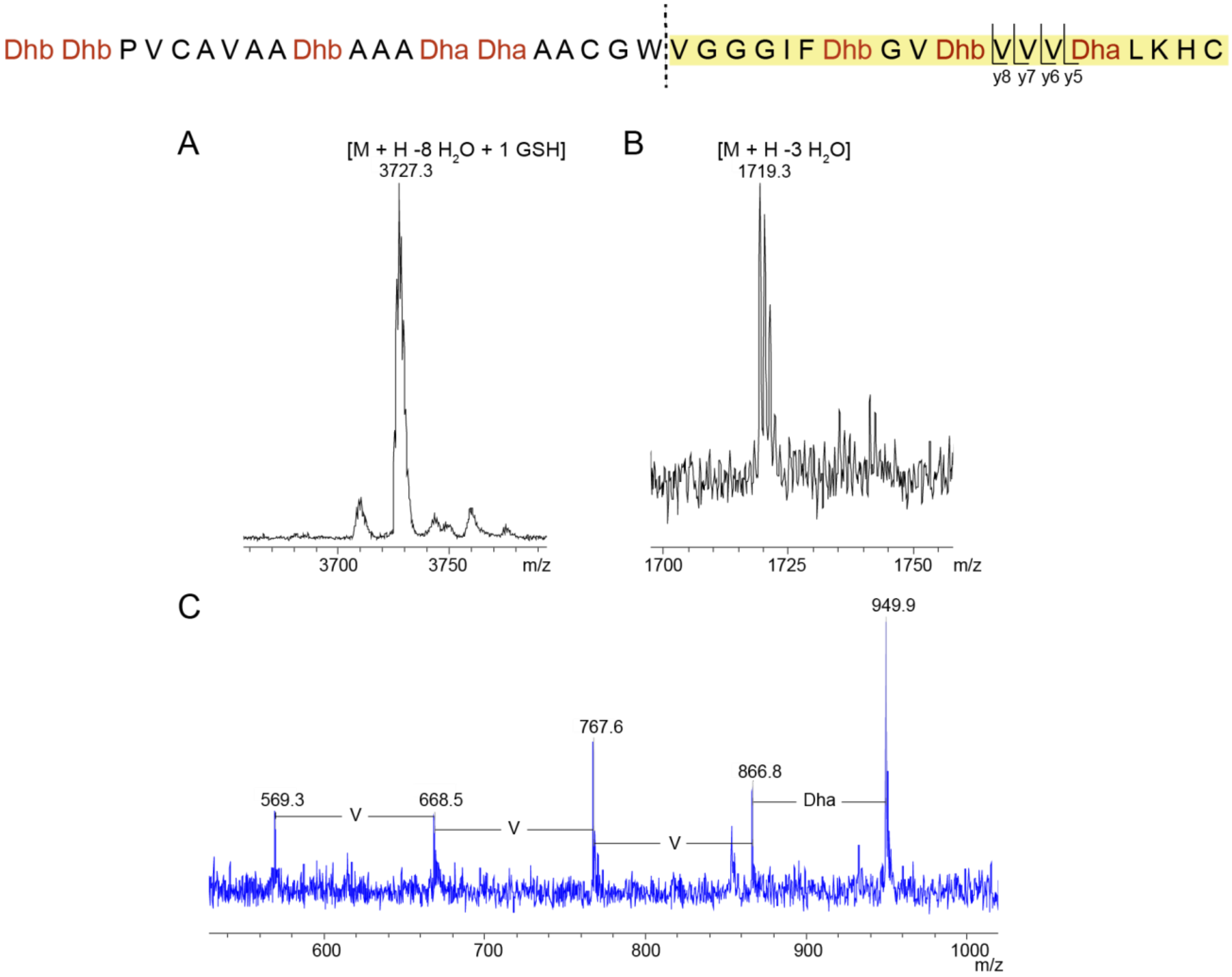


Figure S2: Presence of glutathione adduct in CylL$_L$ co-expressed with CylM in HEK293 cells. A) MALDI-TOF mass spectrum of 8-fold dehydrated CylL$_L$" with a GSH adduct. The peptide was purified via Ni-NTA affinity chromatography and digested with CylA. B) MALDI-TOF mass spectrum of the yellow highlighted CylL$_L$ fragment post-chymotrypsin digest. C) LIFT analysis of the 1719 Da fragment.

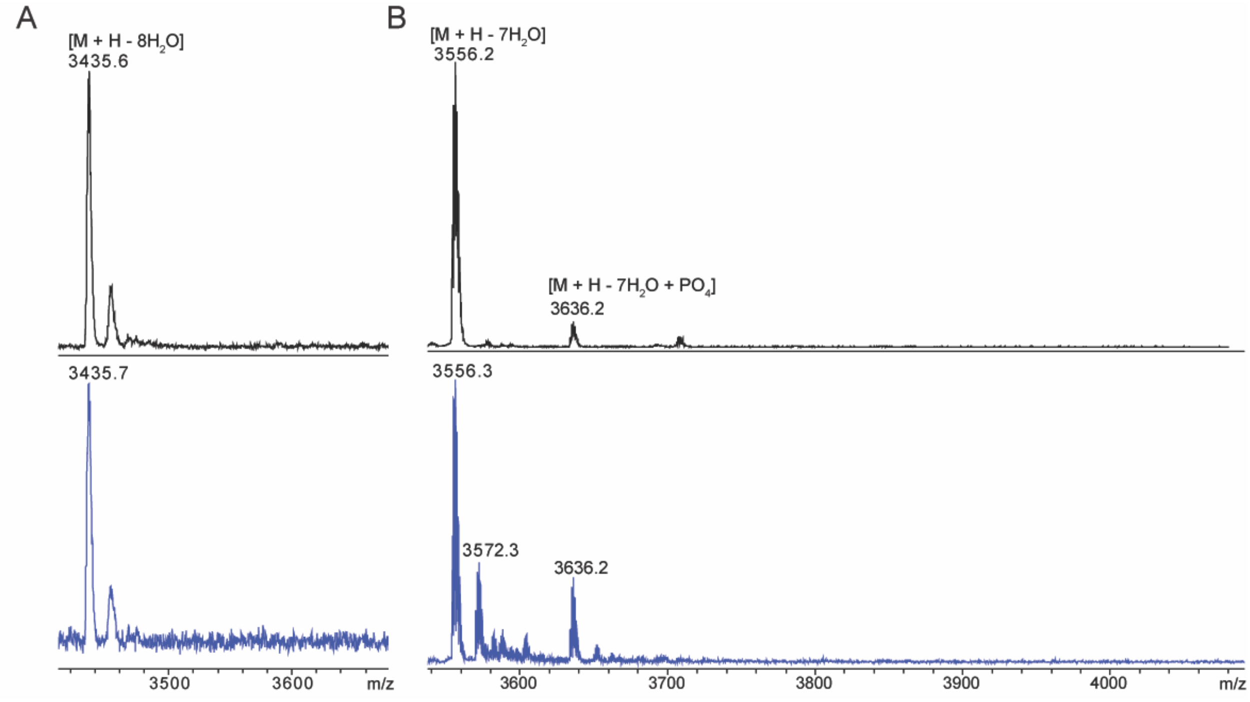


Figure S3: MALDI-TOF mass spectra of (A) CylL$_L$-S15T and (B) HalA2 co-expressed with CylM and HalM2, respectively, before (black) and after (blue) IAA (A) or NEM (B) reaction. Peptides were expressed in Expi293 cells and purified via Ni-NTA chromatography and analytical HPLC. CylL$_L$-S15T and HalA2 were digested with CylA and GluC, respectively. Reaction products were desalted via C4 ziptip.

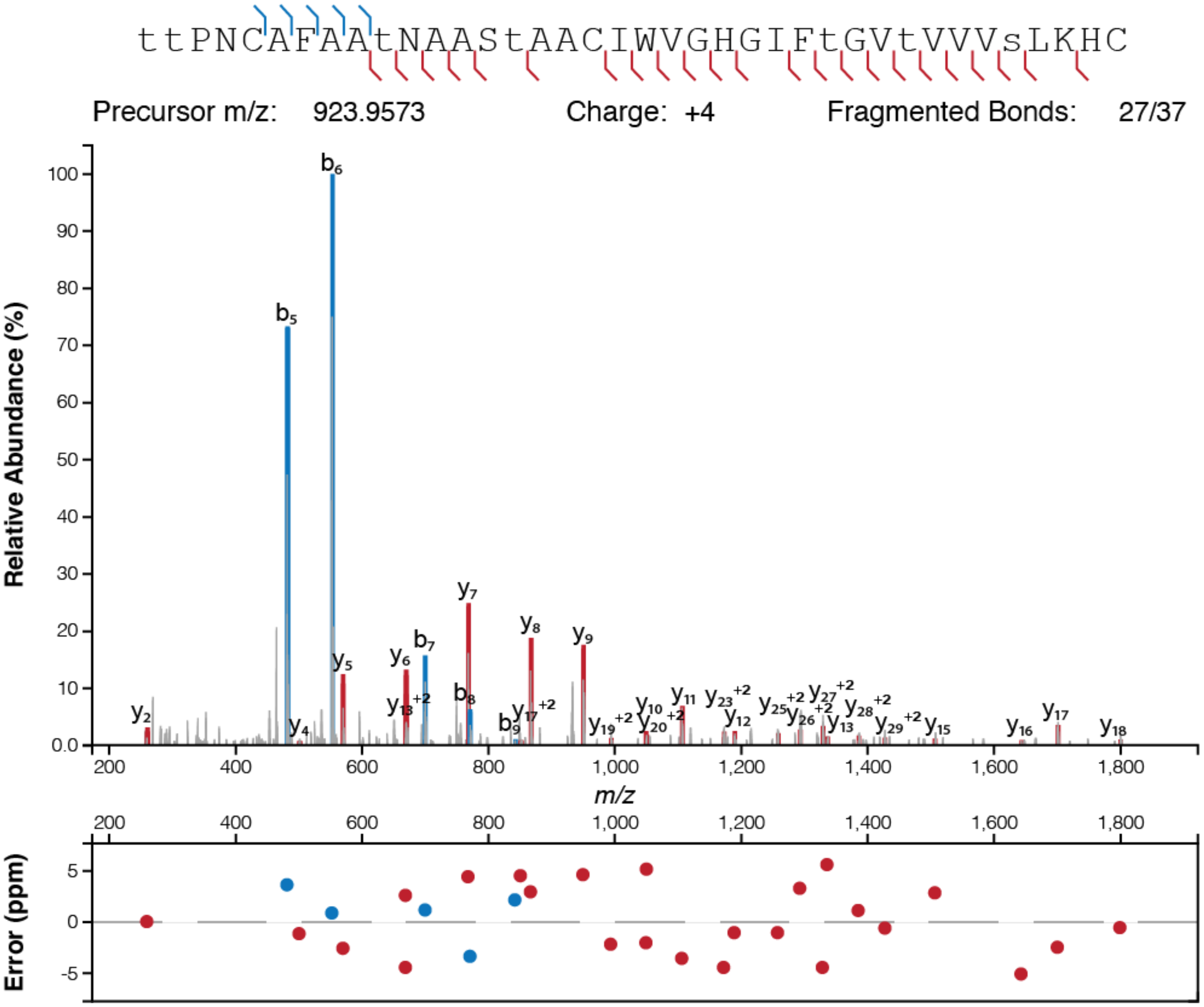


Figure S4: High-resolution MS/MS spectrum of CylL$_L$-S15T variant NDT1 co-expressed with CylM in Expi293 cells. Peptide was purified via Ni-NTA affinity chromatography and digested with CylA. A graph of the ppm errors for each identified ion is shown.[74]

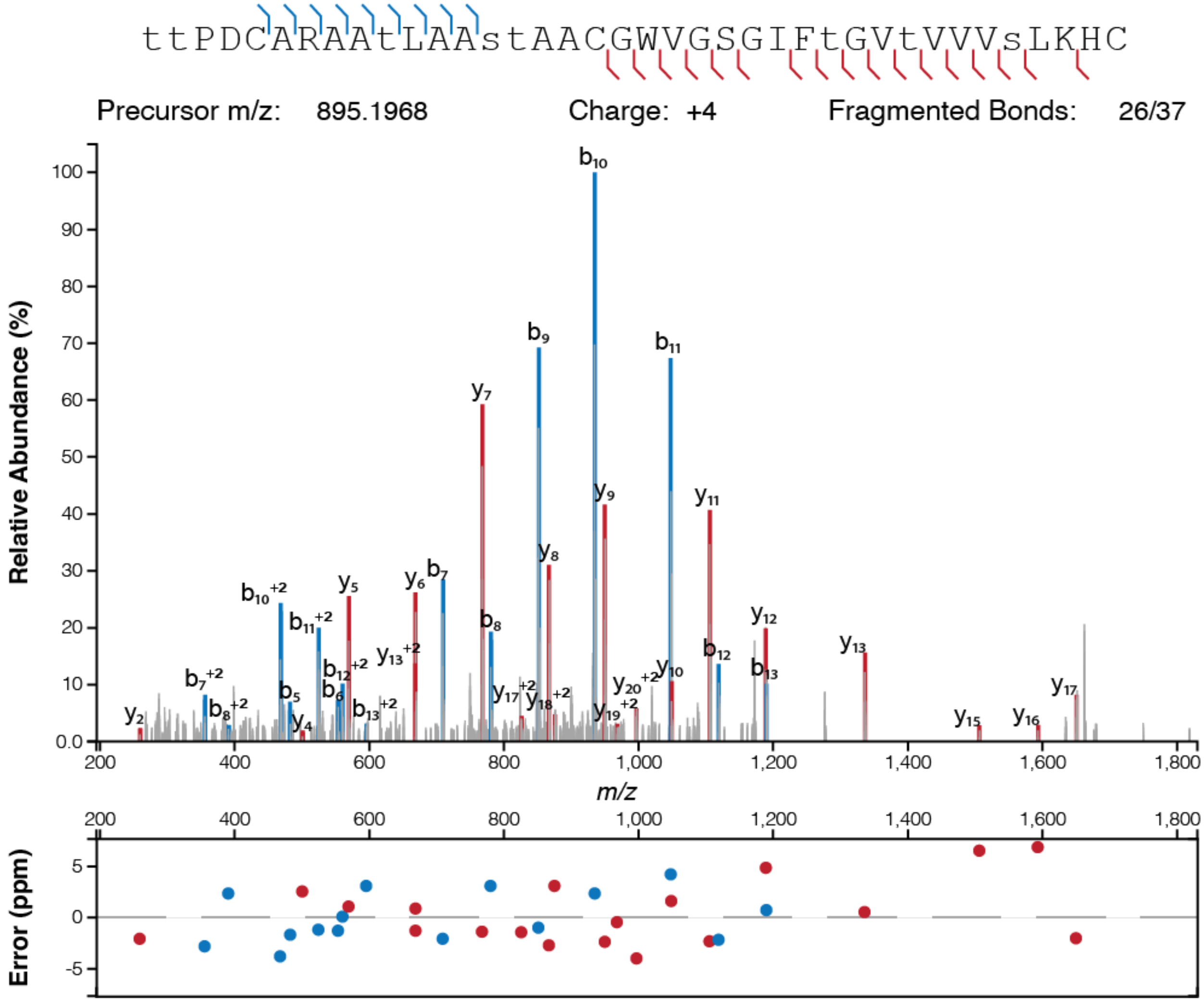


Figure S5: High-resolution MS/MS spectrum of CylL<sub>L</sub>-S15T variant NDT2 co-expressed with CylM in Expi293 cells. Peptide was purified via Ni-NTA affinity chromatography and digested with CylA. A graph of the ppm errors for each identified ion is shown.[74]

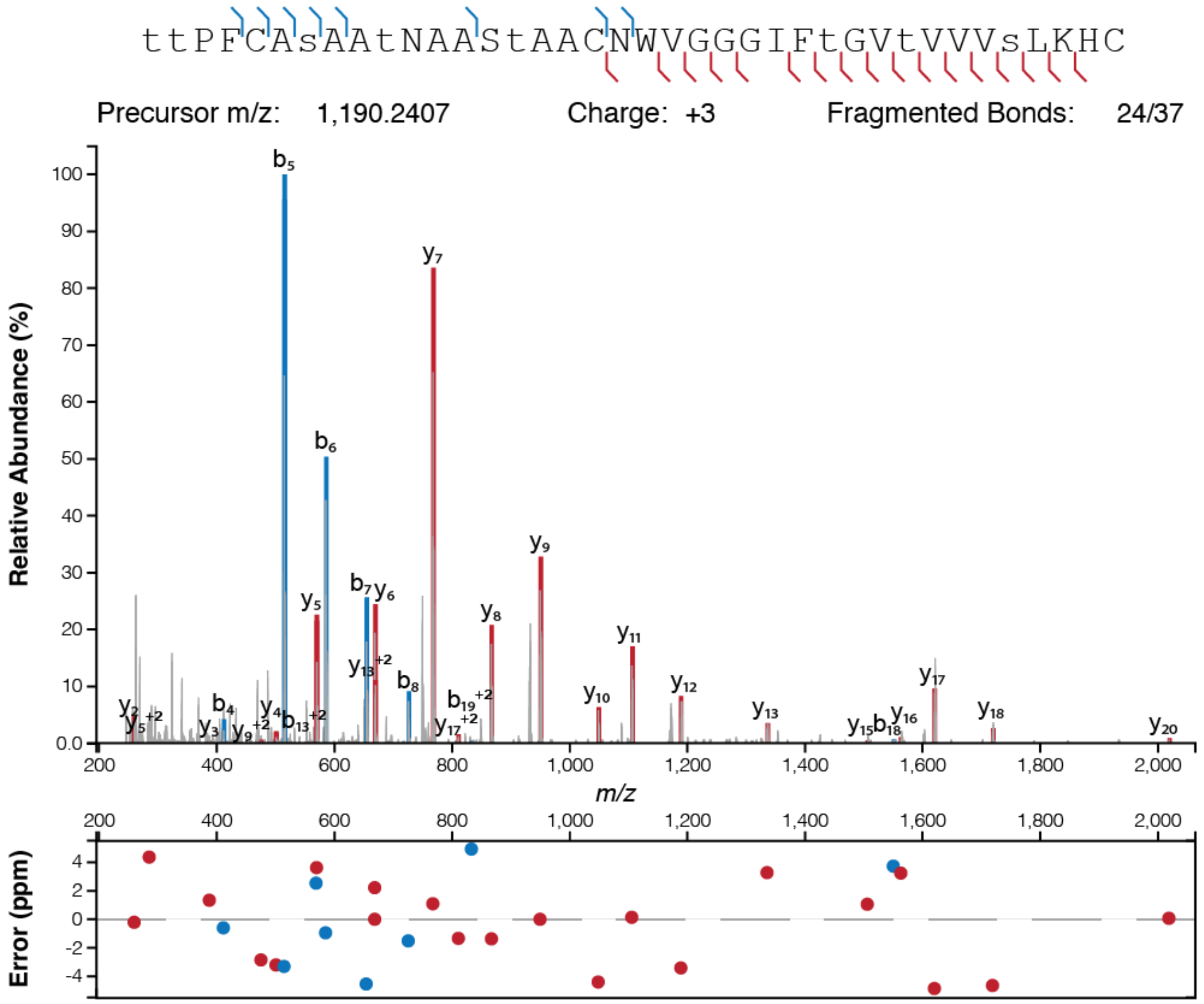


Figure S6: High-resolution MS/MS spectrum of $\text{CylL}_{\text{L}}$”-S15T mutant NDT3 expressed in Expi293. Peptide was purified via Ni-NTA affinity chromatography and digested with CylA. A graph of the ppm errors for each identified ion is shown.[74]

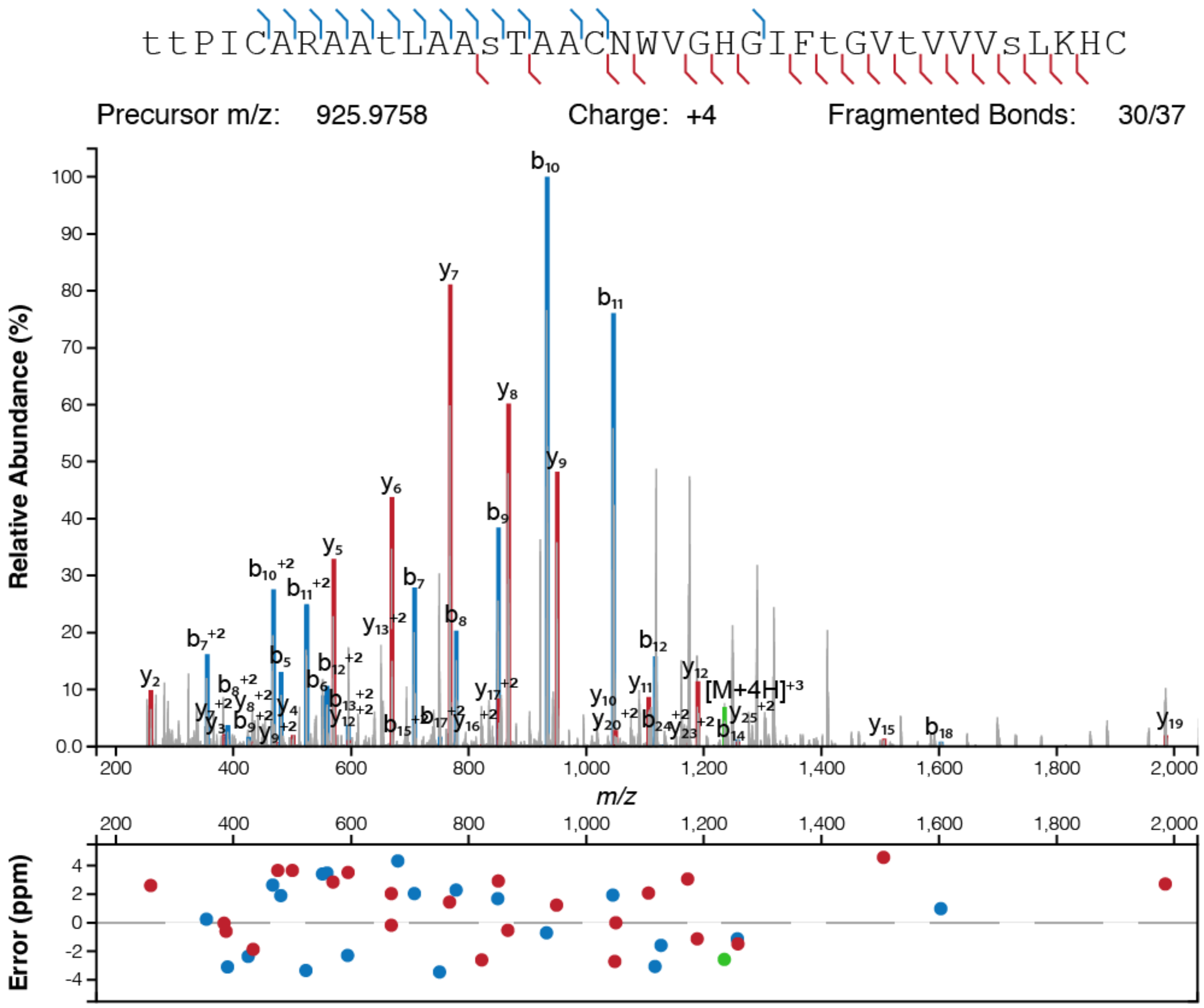


Figure S7: High-resolution MS/MS spectrum of CylL$_L$”-S15T mutant NDT4 expressed in Expi293. Peptide was purified via Ni-NTA affinity chromatography and digested with CylA. A graph of the ppm errors for each identified ion is shown.[74]

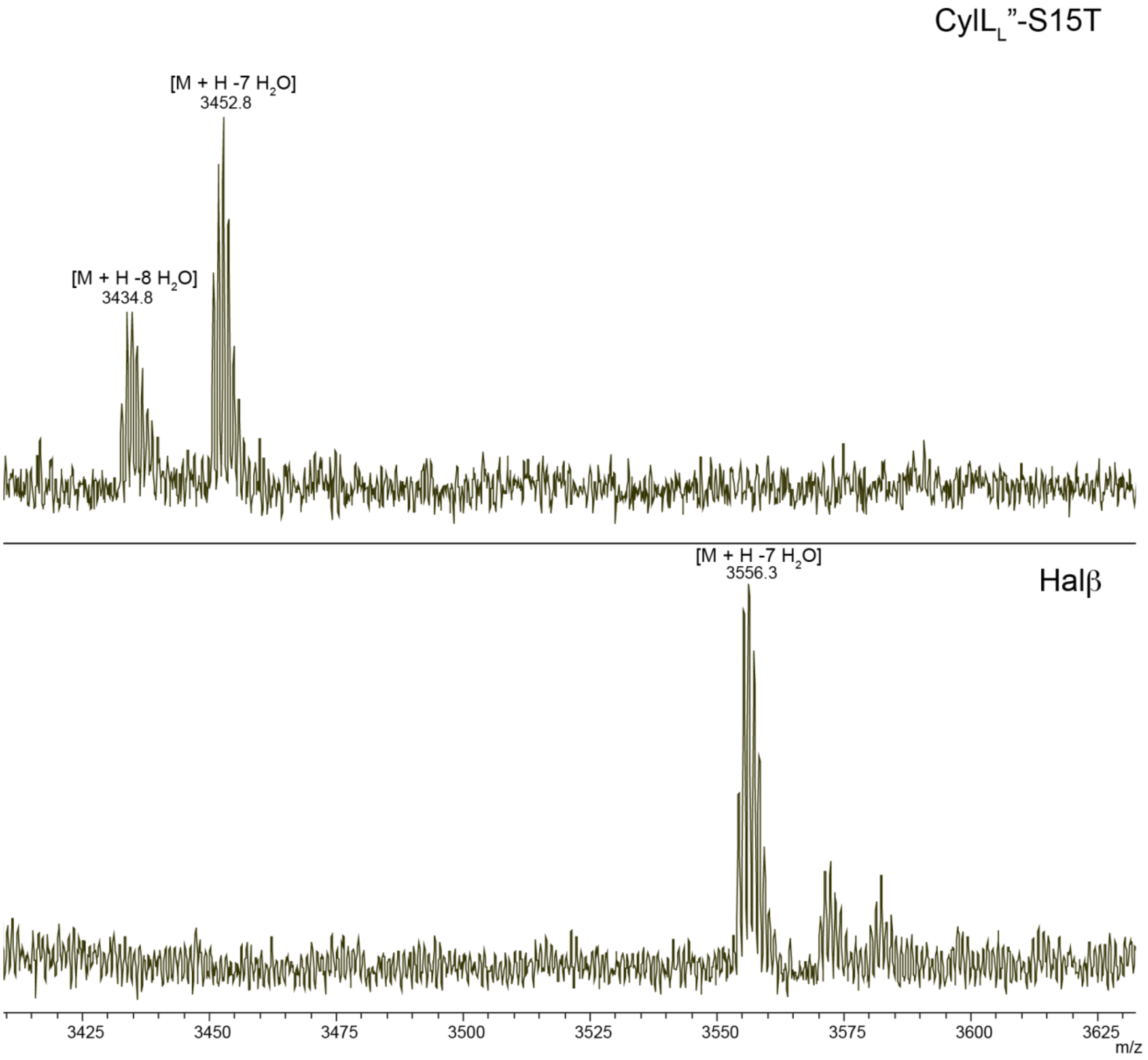


Figure S8: MALDI-TOF mass spectra of nuclear targeted CylL$_L$-S15T and nuclear targeted HalA2 co-expressed with CylM and HalM2, respectively, in Expi293 cells. CylL$_L$-S15T and HalA2 were digested with CylA and GluC respectively.
